## Supplementary material for "SARS-CoV-2 within-host population expansion, diversification and adaptation in zoo tigers, lions and hyenas": Table S1, Figure S1 and GISAID acknowledgments

Sue VandeWoude

**Supplementary Tables:**

**Table S1.** Available below.

Available in DZ_supplementary.xlsx file:

**Table S2.** All mutations detected relative to Wuhan reference sequence, including AY20 characteristic mutations and within-host mutations (relative to tiger reference).

**Table S3.** Characteristic AY.20 mutations from outbreak.info.

**Table S4.** MultiQC summary containing coverage and quality metrics output by the nf-core/viralrecon pipeline.

**Table S5.** Variant table output by the nf-core/viralrecon pipeline.

**Table S6.** Population-level nucleotide diversity measures for full SARS-CoV-2 genomes.

**Table S7.** Gene-level nucleotide diversity measures.

**Supp. Information:** GISAID acknowledgements available below.

**Figure S1.** Available below.

**Table S1**. **Denver Zoo animal metadata reproduced with permission from Gallichotte et al. (2024).**

| **Species** | **Group** | **Animal ID** | **Sex** | **Age*** |
| --- | --- | --- | --- | --- |
| Amur tiger | --- | A | F | 11 |
|  |  | B | M | 11 |
| African lion | pride 1 | A | M | 6 |
|  |  | B | M | 6 |
|  |  | C | M | 6 |
|  |  | D | M | 6 |
|  | pride 2 | E | F | 6 |
|  |  | F | F | 1 |
|  |  | G | M | 5 |
|  |  | H | F | 9 |
|  |  | I | M | 1 |
|  |  | J | F | 9 |
|  |  | K | M | 2 |
| Spotted hyena | older clan | A | F | 22 |
|  |  | B | M | 23 |
|  | young clan | C | M | 7 |
|  |  | D | F | 7 |

*Age at the time of the outbreak (Oct-Nov 2021).

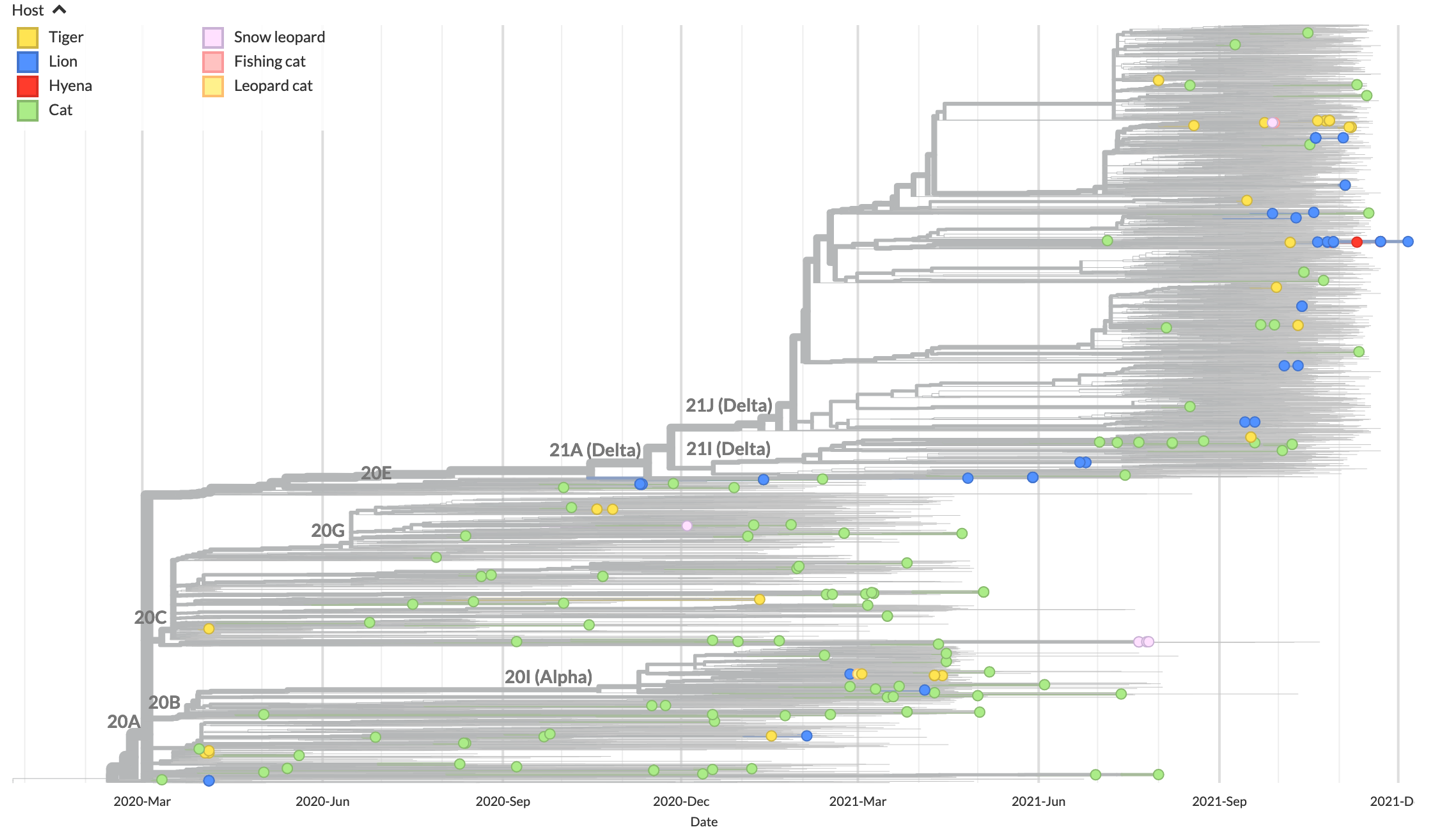

**Figure S1.** No evidence at the global level for species-specific adaptation of SARS-CoV-2 in felids or hyenas. A time-based phylogenetic tree was generated from from all felid- and hyena-derived SARS-CoV-2 sequences available in the GISAID database prior to December 2021, with contextual human-derived sequences collected in the same time and place. Tree was generated with Nextstrain and an interactive version is available at <https://nextstrain.org/community/laurabashor/DZSARS2>. Sequences are colored by host species and scaled by time, with collection date indicated along the x-axis. four data inputs: (1) sixteen SARS-CoV-2 consensus sequences from the Denver Zoo outbreak corresponding to the highest quality sequence collected at the latest date from each individual animal, (2) a random subsample of human-derived sequences in Colorado from the days leading up to the zoo outbreak (100 sequences/day from September 23rd to October 7th, 2021; N=1500 sequences), (3) all felid-derived SARS-CoV-2 sequences available in the GISAID database prior to December 2021, and (4) contextual sequences collected in the same time and place as each felid sequence (10-100 human-derived sequence per felid-derived sequence) with enriched sampling within the United States. All data obtained from GISAID were processed with the Augur pipeline for input into Nextstrain, and acknowledgements are available in GISAID EPI_SET_240407wv.

**Sequences from GISAID included in this study**

We prepared two GISAID EPI_SETs to acknowledge publicly available SARS-CoV-2 sequences included in this study. EPI_SET_240407tz includes all sequences used to generate the phylogenetic tree in Figure 2A, and additional individual GISAID sequences discussed in the text. EPI_SET_240407wv includes all additional sequences used to generate the phylogenetic tree in Supplementary Figure 1.

**Data Availability**

GISAID Identifier: EPI_SET_240407tz
doi: 10.55876/gis8.240407tz

All genome sequences and associated metadata in this dataset are published in GISAID’s EpiCoV database. To view the contributors of each individual sequence with details such as accession number, Virus name, Collection date, Originating Lab and Submitting Lab and the list of Authors, visit 10.55876/gis8.240407tz

**Data Snapshot**

EPI_SET_240407tz is composed of 1,525 individual genome sequences.
The collection dates range from 2019-12-31 to 2021-12-21;
Data were collected in 4 countries and territories;
All sequences in this dataset are compared relative to hCoV-19/Wuhan/WIV04/2019 (WIV04), the official reference sequence employed by GISAID (EPI_ISL_402124). Learn more at https://gisaid.org/WIV04.

**Data Availability**

GISAID Identifier: EPI_SET_240407wv
doi: 10.55876/gis8.240407wv

All genome sequences and associated metadata in this dataset are published in GISAID’s EpiCoV database. To view the contributors of each individual sequence with details such as accession number, Virus name, Collection date, Originating Lab and Submitting Lab and the list of Authors, visit 10.55876/gis8.240407wv

**Data Snapshot**

EPI_SET_240407wv is composed of 11,999 individual genome sequences.
The collection dates range from 2019-12-31 to 2021-11-22;
Data were collected in 26 countries and territories;
All sequences in this dataset are compared relative to hCoV-19/Wuhan/WIV04/2019 (WIV04), the official reference sequence employed by GISAID (EPI_ISL_402124). Learn more at https://gisaid.org/WIV04.
